# Fine-scale coexistence between Mediterranean mesocarnivores is mediated by spatial, temporal and trophic resource partitioning

**DOI:** 10.1101/2020.11.05.370403

**Authors:** Iago Ferreiro-Arias, Jorge Isla, Pedro Jordano, Ana Benítez-López

## Abstract

1. The partition of the ecological niche can enhance the coexistence of predators due to differences in how they exploit three main resources: food, space, and time, the latter being an axis that often remains unexplored.
2. We studied niche segregation in a Mediterranean mesocarnivore community composed by *Vulpes vulpes, Genetta genetta, Meles meles* and *Herpestes ichneumon*, addressing simultaneously different niche axes: the temporal, trophic and spatial axes.
3. We assessed temporal segregation between mesopredators and prey and between potential competitors, using camera trap data between 2018-2020 in a Mediterranean landscape in Southern Spain. We deployed camera traps in 35 stations in three sites with varying vegetation cover within Doñana National Park. We further examined the spatial overlap in activity centers and trophic preferences between potential competitors using diet information from studies performed in the study area.
4. We found an overall temporal segregation between trophic generalist species, with species showing higher temporal overlap differing in their trophic preferences and/or showing limited spatial overlap. Furthermore, we observed an overall high overlap between the activity patterns of predators and their major prey in the area (the common genet vs small mammals and the red fox vs European rabbit).
5. Our study suggests that coexistence of the different species that compose the mesocarnivore assemblage in Mediterranean landscapes can be facilitated by subtle differences along the three main niche axes, with temporal segregation being a most pronounced mechanism. Our findings reinforce the idea that the coexistence mechanisms underlying community structure are multidimensional.

## Introduction

Understanding the mechanisms that promote coexistence of species with similar ecological requirements is a central topic in community ecology, where competitive interactions between species have the potential to affect diversity patterns by limiting or promoting coexistence (Chesson, 2000). In order to mitigate the negative impact of interspecific competition, species often partition resources along the temporal, trophic and spatial dimensions, which eventually result in niche differentiation (Schoener, 1974). Most studies concerning species coexistence focus on the differential use of habitat and food resources; nonetheless, the differential use of the diel cycle may enhance coexistence of same-sized species, particularly among predator species (Díaz□Ruiz et. al., 2009; Wang & Fisher, 2012).

A large variability and plasticity in activity patterns has been documented for mammals (Bennie et al. 2014) being mainly determined by circadian endogenous boundaries (Kronfeld-Schor & Dayan, 2003), but also shaped by external factors (Monterroso et al., 2014; Zielinski, 1986). This diversity of diel cycles could be due to the plastic nature of behavioural responses to different pressures, which may in turn induce marked variations in daily rhythms among different scenarios (Ensing et al., 2014; Gaynor et. al., 2018). For example, predators can adjust their daily activity rhythms to those hours of the day with increased availability of prey (Foster et al., 2013), while, at the same time, prey species could generate a temporal mismatch by centering their activity at times when there is a lower risk of predation (Fenn & Macdonald, 1995). Consequently, high temporal overlap between carnivores and their putative prey have been reported in some predator-prey systems (Linkie & Ridout, 2011), whereas in others, asynchronous activity peaks has been reported (Arias-Del Razo et al., 2011; Díaz□Ruiz et al., 2016).

Moreover, the co-occurrence of species that compete for a certain resource has the potential to produce shifts in the daily activity patterns of competitively inferior species. For example, pampas foxes have been reported to change their daily foraging activity in areas where the dominant fox species, the crab-eating fox, is present. In areas where only pampas foxes are present, the species shows a nocturnal pattern whereas in sympatric areas, pampas foxes switch from nocturnal to diurnal to reduce their activity during periods when the dominant crab-eating fox is active. (Di Bitetti et al., 2009). As a consequence, the daily activity of a given predator could be a result of a cost-benefit trade-off between maximizing their activity during periods with high prey availability (benefit) but also higher mortality risk due to interspecific encounters with intraguild competitors (cost) (Santos et al., 2019).

Interspecific interactions therefore play an important role in structuring behavioural spatio-temporal dynamics within mammalian communities (Holt et al., 1994), especially in carnivores where competition not only arises as a consequence of exploiting the same resource simultaneously, but also due to the risk associated with intraguild killing (IGK). Intraguild killing is an antagonistic interaction where a predator species kills another predator, or where both species prey upon each other (Palomares & Caro, 1999; Polis et al., 1989), being frequently reported among top-predators against their inferior competitors (Palomares & Caro, 1999). The IGK’s effects in competitors can be diverse, from local abundance reductions and distributional shifts in space (Jiménez et al., 2019; Newsome et al., 2017) to behavioural modifications through restriction of their activity patterns (Wang & Fisher, 2012). Because apex predators play a key role in ecosystem functioning due to their regulatory role on populations of prey and medium-sized carnivores (top-down control), their extinction frequently triggers cascading effects that can lead to an eventual increase of the distribution and abundance of medium-sized carnivore populations, a phenomenon coined ‘*mesopredator release*’ by Soulé et al. (1988).

In the Iberian Peninsula, the Iberian lynx (*Lynx pardinus*) exerts its role as a top-predator on the populations of surrogate competitors (Palomares et al., 1998; Palomares et al., 1996). The probability of lynx occurrence is negatively associated with the presence of mesopredators (Monterroso et al., 2020; Palomares et al., 1996), and sizeable reductions in mesocarnivore abundance have been reported after the reintroduction of the Iberian lynx in some areas (Jiménez et al., 2019), thus reinforcing the idea that carnivore coexistence in areas where this top predator is present is supported by spatial structuring. In Doñana National Park, the carnivore guild includes the Iberian lynx, a habitat and trophic specialist top predator, and mesocarnivores such as the red fox (*Vulpes vulpes*), the European badger (*Meles meles*) and the Egyptian mongoose (*Herpestes ichneumon*), which are characterized by a wider habitat and trophic niche breadth. This assemblage is completed by the common genet (*Genetta genetta*), with a considerably narrower diet. The negative effects that the Iberian lynx has on these mesocarnivores mainly emerge through intraguild killing, particularly on Egyptian mongooses (*Egyptian mongoose)*, common genets (*Genetta genetta)* and red foxes (*Vulpes vulpes*) (Palomares et al., 1996; Palomares & Caro, 1999, and references therein). Yet, in Doñana, the Iberian lynx population has undergone a considerable redistribution recently, expanding beyond the limits of the national park (López-Parra et al., 2012), and it is currently absent in some areas where its top-control over the mesopredator assemblage is thus relaxed, and where interspecific competition between mesocarnivores may become more relevant.

Because these generalist mesopredators exhibit no apparent differences in habitat use (Soto & Palomares 2015), in areas with limited space and/or reduced trophic resources coexistence may be achieved by fine-scale segregation of activity patterns. For example, a reduction in the abundance and availability of the main shared trophic resource at a given moment may lead to increased competition for the consumption of available secondary resources. A clear example is the reported reduction in the consumption of European rabbits after a sudden population collapse due to the emergence of the rabbit hemorrhagic disease (RHD) (Ferreras et al., 2011), which resulted in an increased trophic overlap among most generalist carnivore species. In such cases, time partitioning becomes the main mechanism to facilitate coexistence by reducing competitive stress in resource use among these species (Barrientos & Virgós, 2006). Nevertheless, empirical data on fine-scale temporal activity patterns are surprisingly limited for the Mediterranean area, with most studies focusing on spatial segregation at meso-to macro-scales as the main mechanism allowing mesocarnivore coexistence (Fedriani, 1999; Soto & Palomares 2015; Monterroso et al., 2020). While recent studies have focused on temporal partitioning (Barrull et al., 2014; Monterroso et al., 2014; Vilella et al., 2020), the spatial scale of these studies is too broad for capturing potential antagonistic interactions. For example, the potential for subtle temporal shifts to reduce interspecific encounters may be tightly coupled with differences in fine-scale spatial occupancy patterns. Detecting such effects requires repeated sampling and monitoring over reduced spatial extents and extended periods of time.

Here, we evaluate the main mechanisms that allow the coexistence of mesocarnivore species in the Doñana National Park in sites where the Iberian lynx is completely absent. We explored niche segregation along the temporal axis for four mesocarnivore species: the red fox (*Vulpes vulpes*), the European badger (*Meles meles*), the common genet (*Genetta genetta*), and the Egyptian mongoose (*Herpestes ichneumon)* (Figure 1). We also assessed evidence of spatial avoidance over fine spatial scales, to understand if avoidance mechanisms can be detected at the scale that IGK might occur (i.e., as a result of interspecific encounters within a few hundred meters). Finally, we complement our findings with an analysis of trophic requirements based on the literature of diet composition for the four species performed in Doñana National Park. Thus, this study aims: 1) to explore the relationship between mesocarnivores along the trophic niche axis; 2) to determine the diel activity patterns of predator and prey species and their synchrony; 3) to quantify the temporal overlap or segregation between potential competitors; and 4) to quantify the spatial overlap among predators’ activity centers.

**Figure 1.**
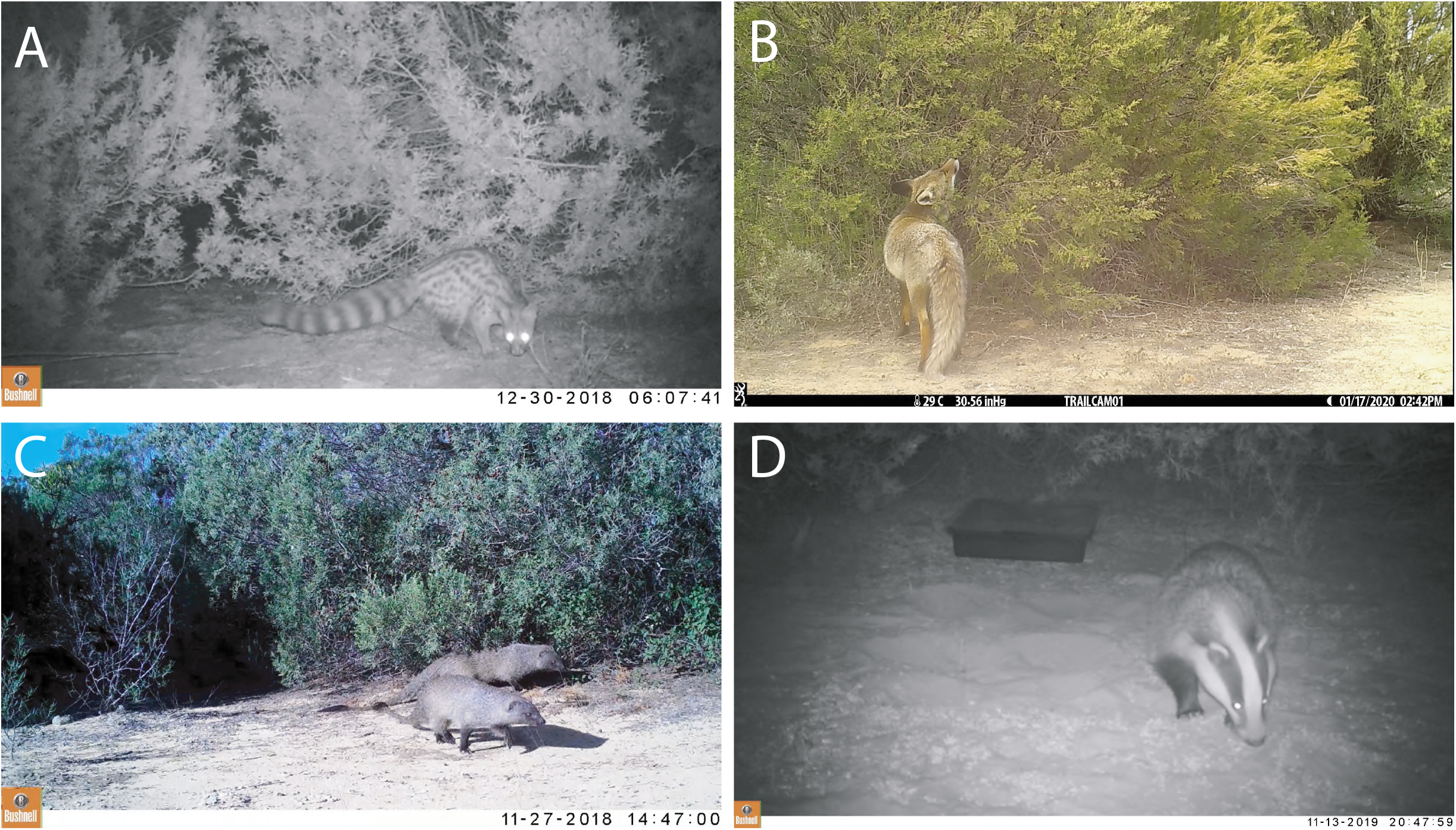
Mesocarnivore target species. From top-left to bottom-right: A, common genet (*Genetta genetta*, 1.2-2.5 kg); B, red fox (*Vulpes vulpes*, 3-14 kg); C, Egyptian mongoose (*Herpestes ichneumon*, 2-2.5 kg) and D, European badger (*Meles meles*, 4.8-9.8 kg).

Following a niche partitioning conceptual framework, we hypothesize a greater temporal segregation among species that have a higher degree of overlap in their respective trophic and/or spatial niches (i.e., between foxes and badgers or mongooses and genets). Conversely, coexistence between species that have similar use of the diel cycle (i.e., genets and badgers) would be mediated by differences in trophic preferences and/or fine-scale spatial avoidance. Furthermore, we expect that predators with narrower trophic spectra would synchronize their activity patterns to those of their preferred prey species (i.e., genets and small rodents), while generalist predators would not show such a marked pattern. Finally, we expect that activity synchrony between generalist predators and prey would be modulated by differences in vegetation structure and cover, and thus, in prey availability (e.g., European rabbit and rodents) among sites.

## Material and methods

### Study area

This study was carried out in Doñana National Park (37°1’N, 6° 33’W), SW Spain. Doñana National Park comprises 54,251 ha endowed with a Mediterranean climate with Atlantic influence, characterized by hot and dry summers and mild and rainy winters (462mm of average annual rainfall). We selected three 1-ha study sites “Sabinar de Colonización” (SABCOL), “Sabinar del Ojillo” (SABOJI) and “Sabinar del Marqués” (SABMAR), within the Doñana Biological Reserve (RBD) (Figure 2). RBD comprises 6,794 ha located in the core area of the Doñana National Park in which three main land cover types can be distinguished: Mediterranean shrubland (80% ca. of RBD’s surface) marshland (12%), and sand dunes (8%). Study sites are characterized by juniper stands (*Juniperus phoenicea subsp. turbinata*) of different age and ecological succession stages, ranging from mature stands with a high vegetation cover to an advanced colonization front of juniper stands with lower vegetation cover. “Sabinar del Marqués” (SABMAR) constitutes the most mature stage with 10520 junipers/ha, “Sabinar del Ojillo” (SABOJI) is at an intermediate stage with 9010 junipers/ha, and “Sabinar de Colonización” (SABCOL) represents the colonization front stage with 2030 junipers/ha. We estimated the percentage of vegetation cover (%) along ten 50 m transects within each study site. Woody vegetation cover was *x*□ ± SD = 24.4 ± 20.6 for SABCOL; 79.1 ± 60.7 for SABOJI and 94.7 ± 97.34 for SABMAR (Figure 2).

**Figure 2.**
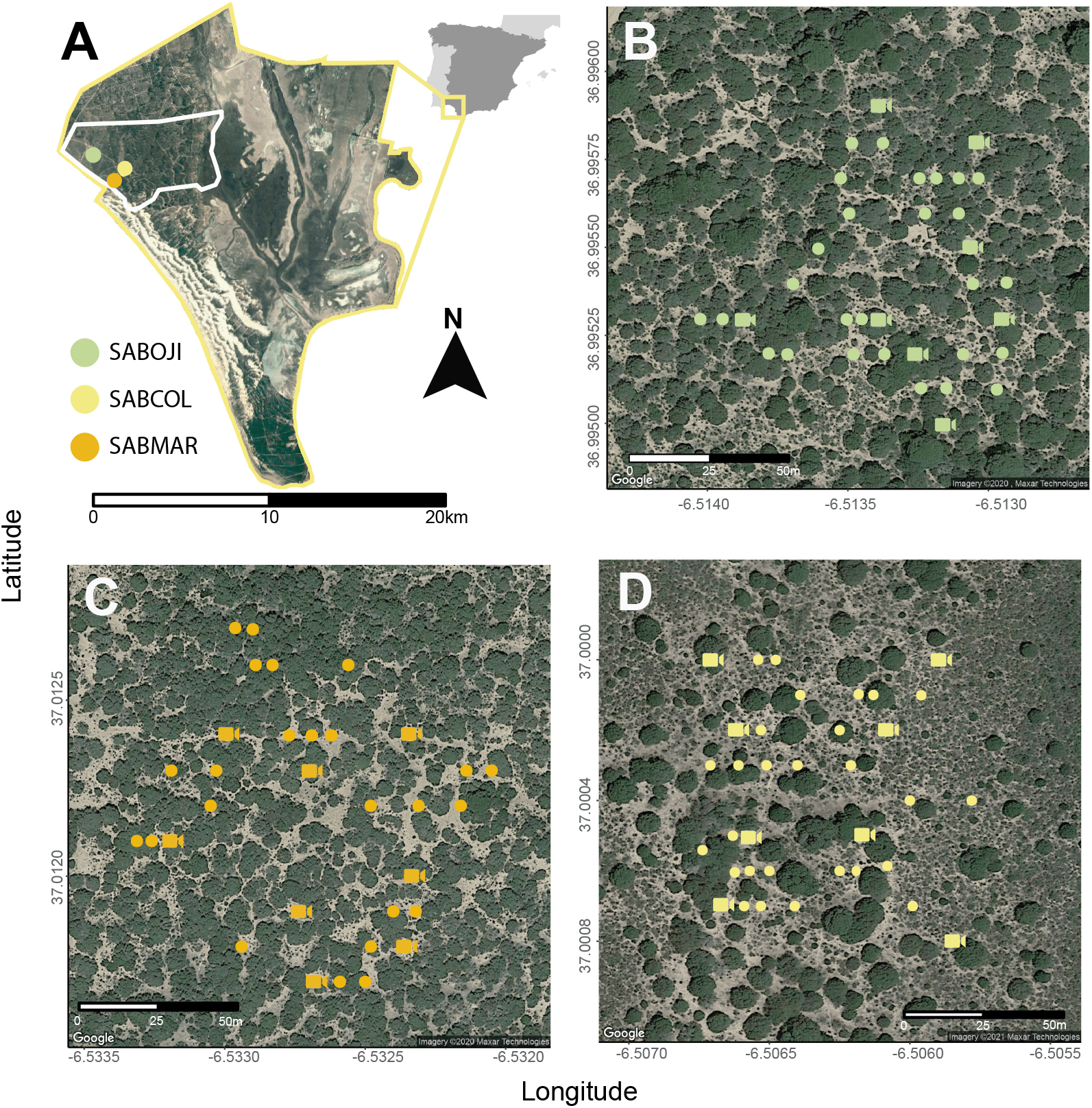
Study area showing: A, location of Doñana National Park and Doñana Biological Reserve (white rectangle) and the location of 3 study sites selected (Sabinar de Colonización - SABCOL, Sabinar del Ojillo - SABOJI and Sabinar del Marqués - SABMAR). B, C and D subplots represent the location of camera trapping stations (points) and set of cameras active for ca. 10 days during a given sampling period within SABOJI, SABMAR and SABCOL respectively. Distances between SABCOL-SABMAR, SABCOL-SABOJI and SABMAR-SABOJI are 0.98, 2.69 and 2.71 km respectively.

### Target species

Our target species consisted of the four mesopredators which occur in Doñana National Park namely the Egyptian mongoose (*Herpestes ichneumon*), the red fox (*Vulpes vulpes*), the European badger (*Meles meles*) and the common genet (*Genetta genetta)* (Valverde, 1967) (Fig. 1), and their respective prey species. Despite being present in specific areas of DNP, the Iberian lynx (*Lynx pardinus*) was not present in our study sites, thus not exerting its role as top predator and influencing spatio-temporal dynamics of the carnivore assemblage. Potential prey species include lagomorphs (mainly European rabbits, *Oryctolagus cuniculus*), small mammals and birds such as the red partridge *Alectoris rufa*. In Doñana, the European rabbit accounts for a very high percentage of the diet of red foxes (Rau, 1987), Egyptian mongooses (Palomares & Delibes, 1991b) and also for European badgers. Some authors even pointed out a possible specialization of badgers for this type of prey (Martín et al., 1995), an assertion which has been refuted (Revilla & Palomares, 2002). Small mammals and birds are preyed mostly by the common genet, although they are also widely consumed by the Egyptian mongoose (Palomares & Delibes, 1991a) and, to a lesser extent, by the red fox and European badger. Detections from ungulates (mainly *Cervus elaphus* and *Sus scrofa*) were not included in the analysis as their presence in mesocarnivore diets is frequently associated with carrion consumption, with direct predation being recorded sporadically on very young individuals (Valverde, 1967). Conversely, we did not have sufficient detections from the other lagomorph present in RBD (*Lepus granatensis*, N = 3) to include them in this study.

### Diet data

We searched on Web of Science (WOS) and Google Scholar for studies that reported trophic preferences of the four mesocarnivores in Doñana National Park using the following search string: (trophic OR diet) AND (predator species) AND (Doñana), where “predator species” was substituted by one of the mesopredator species considered in this study when performing the search. We also gathered diet data for the common genet by consulting experts (P. Ferreras pers. comm.). The selection of studies contains all the trophic information available for the four species at different locations in the Doñana National Park in different years and seasons, thus reflecting the entire trophic spectra of each of the different species (Supplementary Table S1). We collected all food items per species and study without pooling seasons or periods in order to capture the natural variability in dietary composition across the whole Doñana study area along the annual cycle, thus matching the temporal span of our camera trap data (Supplementary Fig. 1.)

Some studies reported information collected in previous studies (e.g., Martín et al. (1995) and Fedriani et al. (1998), or Palomares and Delibes (1991b) and Palomares (1993)). To avoid pseudo-replication issues, we kept only those studies where the data strictly belong to different samples and periods. From each study, period and species, we extracted the frequency of occurrence of food items in mesopredator faeces. In order to estimate trophic preferences, diet breadth and diet similarity between mesocarnivores, we carried out a correspondence analysis (CA) using the frequency of occurrence of each prey item for each predator and employing the “*FactoMineR*” package (Lê, Josse, & Husson, 2008). The CA ordered the different diet studies (cases) relative to the prey species composition reported by each study, resulting in an ordination of the studies corresponding to each mesopredator species based on their similarity of prey species composition. We used the coordinates on the first two CA axes corresponding to each diet study to build convex polygons spanning each mesopredator group of studies. Convex polygons illustrate the range of dietary variation shown by each mesopredator species in relation to the diet composition of other species.

### Camera trap data collection

We deployed eight camera traps per site. In each site, the eight cameras were rotated periodically through 35 randomly selected focal juniper plants (stations) in order to cover all the surface of each study site. Juniper plants were selected employing a stratified random sampling: stations were selected at random in each of five subplots within each of the three main sites, with the subplots distributed regularly throughout the main plot. Mean distance among neighboring selected juniper plants was 48 m for SABCOL, 49 m for SABOJI and 58 m for SABMAR. Each site contains paths of variable width between juniper individuals, where the cameras were installed at 1-m height focusing on the ground and the bottom of the plant. Camera trap models employed in this study were Browning Dark Ops Pro XD and Bushnell Aggressor. Camera traps were active for ca. 10 days in each of these stations and then changed to other 8 locations, thus ensuring that the whole site was sampled in 5 weeks. Following this procedure, we obtained data from a total set of 35 camera trap stations sampling the three sites during two campaigns: October 2018 - May 2019 and October 2019 - June 2020. All sites had a similar sampling effort (mean number of days that cameras were active): *x*□ ± SD = 80.3 ± 28.3 d for SABCOL; 80.4 ± 27.3 d for SABOJI and 86.8 ± 26.0 d for SABMAR. No attractants were used over the two years of sampling.

### Daily activity patterns and temporal overlap

The diel activity cycle of each species was characterized by pooling the total number of detections across all cameras during the whole study period. In order to avoid data dependency, we discarded consecutive detections of a given species within a site (Davis, Kelly, & Stauffer, 2011; Monterroso et al., 2014). When multiple photographs of the same species were taken within a 15-min interval, we considered them as a single capture event to ensure capture independence. Since the area covered by our study sites is limited (ca. 1 ha/site), we deem 15 min a reasonable time period for a species to abandon the site. In cases where a given individual was detected repeatedly without leaving the camera detection zone, only the time of its first detection was considered.

In our analyses we used either the total, absolute number of records per species (e.g., to estimate daily activity patterns of target species) or the relative frequency of records (Relative Abundance Index, RAI). We calculated the RAI for each mesocarnivore in each camera trap station as the number of detections per 100 camera-days for each study site. We extracted time and date from each detection of every camera trap in order to estimate daily activity patterns and temporal overlap between mesocarnivores and their potential prey, and between mesocarnivore potential competitors. As time and dates are variables with a circular nature, diel activity cycles were estimated with the package “*overlap*” (Ridout & Linkie, 2009) using non-parametric Kernel density plots, which provide a density function of the daily activity pattern of a given species (Rowcliffe et al., 2014). Additionally, we classified the daily activity pattern of each species following van Schaik and Griffiths (1996). A species was classified as: diurnal (> 90% of the detections occurring during the daylight), nocturnal (> 90% of the detections obtained at night), crepuscular (> 90% of their detections occur at dawn or dusk) or cathemeral if their activity pattern is distributed uniformly throughout the daily cycle.

We performed pairwise comparisons of the activity patterns of predators, and of predators and prey in each study site only in those cases where we had more than 10 detections (Fisher, 1995). As the present study has been carried out over two years, daylight can vary from one detection to another and induce bias to our results. Species could adjust their daily activity levels depending on the day length, thus not taking this aspect into account when dealing with strictly diurnal or nocturnal species could lead to the misinterpretation of the results and the underestimation of peaks of daily activity (Vazquez et al., 2019). Therefore, we carried out a double average anchoring time transformation to take into account the mean average sunrise and sunset times for the study area (Vazquez et al., 2019) using the “*activity*” package (Rowcliffe, 2019).

We evaluated the extent of temporal segregation between mesopredators using the overlap coefficient (Δ) described by Ridout and Linkie (2009). This coefficient ranges from 0 to 1, where 0 values represent completely different diel activity patterns while 1 represents the maximum overlap between both species. Following Ridout and Linkie (2009), we used Δ_1_ for cases with a sample size lower than 75 detections, and Δ_4_ when sample size was equal or greater than 75 detections. Once the overlap coefficients between the different species were estimated, 99% confidence intervals were calculated through smoothed bootstrap analysis with 10,000 replicates. Overlap coefficients (Δ) and their respective confidence intervals were calculated with the package “*overlap*” (Ridout & Linkie, 2009). Moreover, we evaluated differences in pairwise comparisons of daily activity patterns using the non-parametric Mardia-Watson-Wheeler test (MWW, Batschelet, 1981) using the “*circular*” package (Agostinelli & Lund, 2017). Further, we used percentiles to establish threshold levels of overlap. Thus, Δ> 75th percentile indicates a high temporal overlap between both species, Δ ≤ 50th percentile a low degree of overlap and, finally, intermediate values of these percentiles, 50th <Δ ≤ 75th, denote moderate levels of overlap (Monterroso et al. 2014).

### Spatial distribution overlap

We calculated the pairwise species spatial overlap between the activity centers of mesopredators using Pianka’s O index (PI, Pianka 1973), based on the RAI data for each site. We calculated bootstrapped confidence intervals around PI using the “*spaa*” package (Zhang et al. 2013). The activity centers were spatially represented for each mesocarnivore species and site using the packages “*ggmap*” and “*ggplot2*” (Kahle & Wickham, 2013; Wickham, 2016). All analyses were conducted in R Studio 1.2.5033 Statistical Software (R Core Team, 2020).

## Results

### Trophic segregation among mesocarnivores

The first two dimensions of the correspondence analysis explained 62% of the variability of mesocarnivore diets (Fig. 3). Axis 1, which explained 34.5% of the variation, discriminates the trophic preferences of the European badger and the common genet due to their eminently omnivorous and carnivorous diets, respectively, but not from the rest of the mesocarnivores. The red fox and the European badger showed very close values due to their frugivorous behaviour and the predominant consumption of lagomorphs and invertebrates. Meanwhile, the Egyptian mongoose had the wider trophic preferences, with an intermediate position between the omnivorous diets of the European badger and the red fox, and the mostly-carnivorous diet of the common genet. Simultaneously, axis 2, which explained 27.3% of the trophic variation, is generated by variation between the diets of the red fox and the common genet, with the latter preferring small mammals followed by birds as main food items. However, the second dimension of the correspondence analysis does not discern between the trophic niches of the red fox, the European badger and the Egyptian mongoose, indicating that there is a greater trophic overlap between these three species with respect to the common genet. Further, convex polygons indicate the broader trophic diversification of the generalist species (red fox, European badger and Egyptian mongoose) whose range of prey types tends to be considerably wider compared with the common genet (Fig. 3).

**Figure 3.**
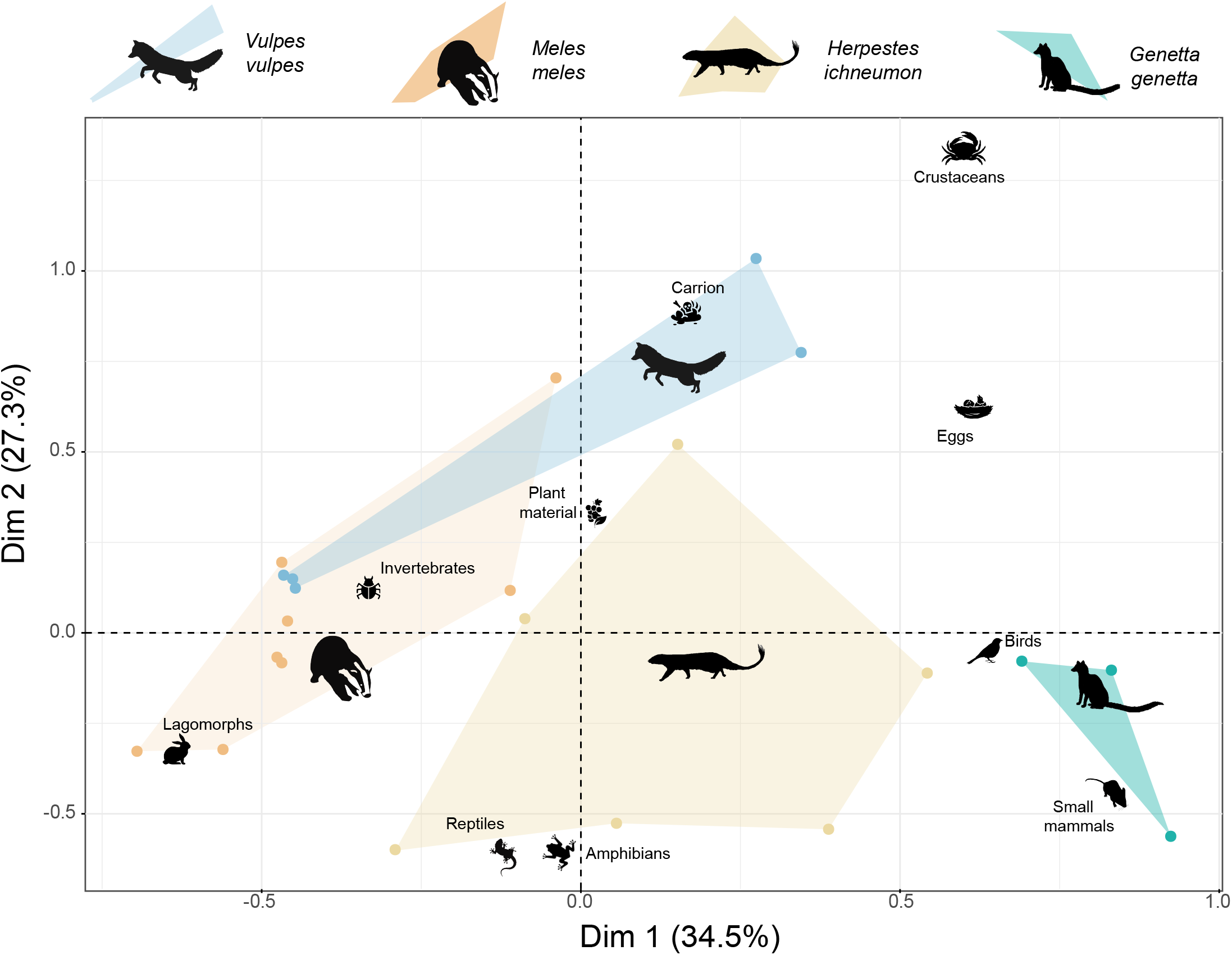
Niche segregation along the trophic axis among mesocarnivores in Doñana National Park. Individual dots indicate species-specific studies of food habits of the four mesocarnivore species. The first two CA axes (Dim 1, Dim 2) explain 66% of the variability in diet composition between species and diet studies. The first dimension represents a gradient of diet variability between the common genet and the European badger, while the second dimension separates the diet of the red fox and the common genet. Convex polygons illustrate trophic niche breadth.

### Species detections and diel activity patterns

We obtained 2786 detections in 531 effective sampling trap-days, but we discarded 96 detections (3.44%) that could not be identified at the species level. Among all valid records, 16% corresponded to mesocarnivores, 28.7% to prey species, 15% to ungulates and 38.7% to passeriformes, which were not included in this study (Supplementary Table S2).

Regarding our target species, we recorded 301 detections of the red fox, with 54.8% of them in SABCOL and the rest of the detections evenly distributed between the other two sites (SABOJI: 23.9%; SABMAR: 21.3%). Most fox detections occurred at night, while the rest were distributed throughout daylight periods, indicating a cathemeral activity pattern with a tendency towards bimodality (Supplementary Table S3). Conversely, detections of European badgers were very scarce in the three study sites (total: 22 detections), with 77% of the detections occurring at night and the rest during the dawn or dusk periods. As for the common genet, its activity pattern was strictly nocturnal (95% of the records produced during nighttime), with most detections occurring in SABCOL (62% of detections). Finally, detections of Egyptian mongooses were mainly diurnal (92% of the detections occurred during daylight), and concentrated in the SABOJI (61%) while the other two sites maintained the same proportion of records (ca. 20%). Due to the small number of detections in each study site, site-level temporal overlap of predators can only be reliably estimated for the red fox and the Egyptian mongoose (Supplementary Table S3).

For prey species, most of the detections belonged to the European rabbit (N = 650 detections). Rabbit detections concentrated similarly between SABOJI and SABMAR, while SABCOL had only 6 detections during the entire sampling period (Supplementary Table S2). Overall, the distribution of rabbit detections spanned throughout the day, with distinct activity peaks around sunrise and sunset (Supplementary Table S3). Small mammals (*Mus spretus* and *Apodemus sylvaticus*) were detected 98 times, mainly in SABMAR and only during nighttime. Finally, red-legged partridges were mostly detected in SABCOL (89% of detections) and showed a markedly diurnal activity pattern (Supplementary Table S3).

### Predator and prey diel activity patterns

Broadly, the activity patterns differed between predators and the potential prey species in the study area. The European rabbit showed a high overlap with the red fox (Δ_4_ = 0.84), followed by a moderate overlap with the European badger (Δ_1_ = 0.62). Conversely, both the common genet and the Egyptian mongoose showed low overlap coefficients (Δ_1_ = 0.47 and Δ_1_ = 0.44, respectively) with rabbit activity patterns (Fig. 4).

**Figure 4.**
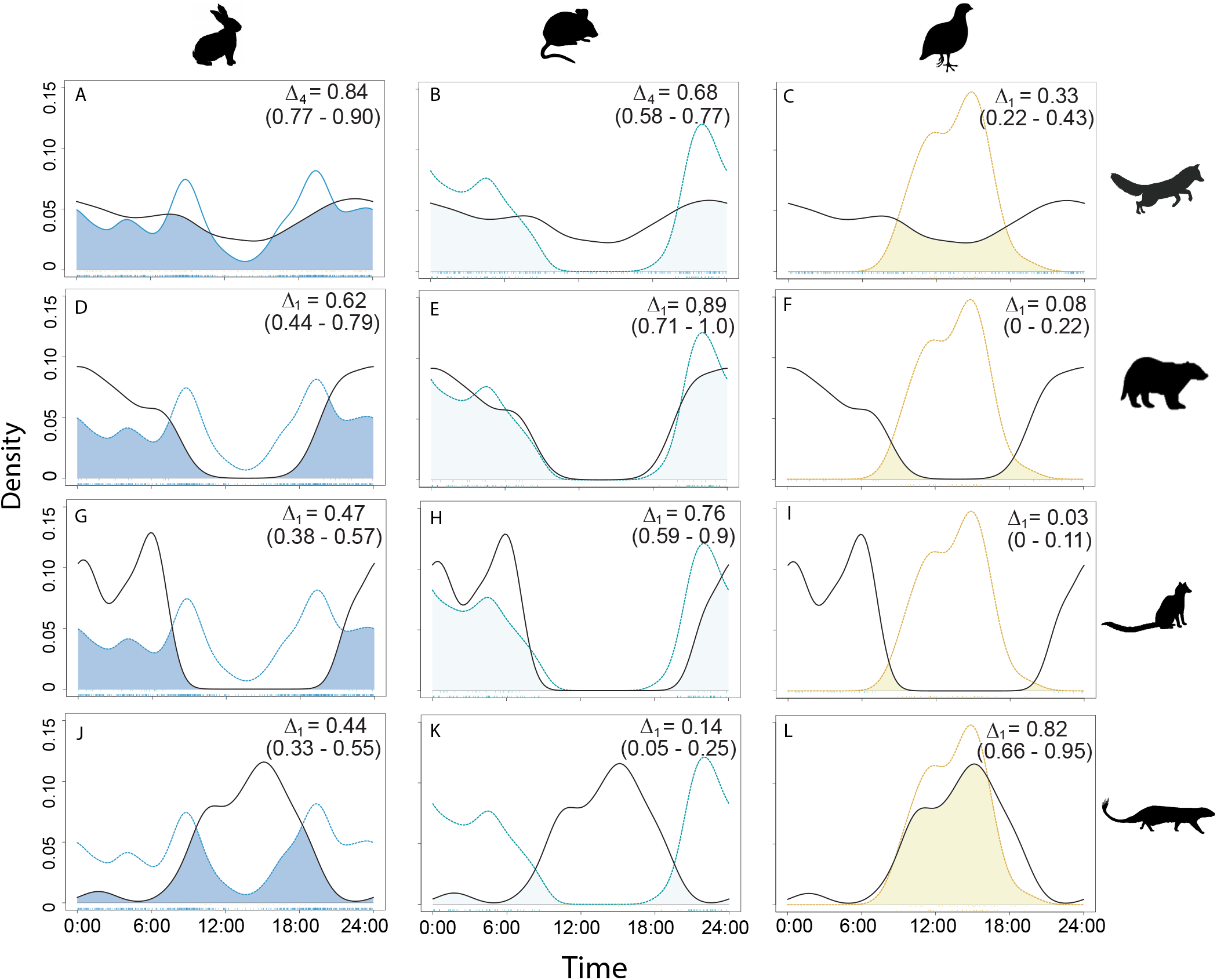
Pairwise comparisons of daily activity patterns between mesocarnivores (rows, top to bottom: red fox, badger, genet, and Egyptian mongoose) and their main prey items in the study area (columns, left to right: *Oryctolagus cunniculus*, small mammals, and *Alectoris rufa)*. Colored lines describe prey activity patterns (blue for European rabbit, green for small mammals and yellow for red-legged partridge) and black lines indicate predator’s activity patterns. Overlap coefficients (Δ) and their respective 99% confidence intervals are shown in brackets.

We found a high temporal overlap between the activity patterns of small mammals (with most detections at night) and the European badger (Δ_1_ = 0.89) and the common genet (Δ_1_ = 0.76). In turn, the red fox showed a moderate temporal overlap with small mammals (Δ_4_ = 0.68), while the Egyptian mongoose showed a total temporal segregation with respect to the detected micromammals (Δ_1_ = 0.14). Activity patterns of red-legged partridges overlapped with those of the Egyptian mongoose (Δ_1_ = 0.82), but not with those of European badgers (Δ_1_ = 0.08) and common genets (Δ_1_ = 0.03) (Fig. 4).

European rabbit activity patterns differed significantly between SABOJI and SABMAR (W = 32.69; p-value < 0.01). Meanwhile, red fox, but not the Egyptian mongoose, showed different responses in terms of diel cycle among sites with different rabbit availability. The red fox showed a predominantly nocturnal activity in SABCOL, where rabbits were absent, and a tendency towards bimodal activity patterns in SABOJI and SABMAR. Red fox activity patterns differed significantly between SABCOL and SABMAR (W = 11.14; p-value < 0.01), while no differences were found between SABCOL - SABOJI (W = 3.97; p-value = 0.137) or SABOJI - SABMAR (W =2.44, p-value = 0.295). In turn, no statistically significant differences were found between activity patterns of the Egyptian mongoose between SABCOL - SABOJI (W = 1.09; p-value = 0.58), SABCOL - SABMAR (W = 2.06; p-value = 0.36) and SABOJI - SABMAR (W =0.40; p-value = 0.82). The temporal overlap between the activity patterns of the European rabbit and the red fox was Δ_1_ = 0.79 for SABOJI and Δ_1_ = 0.73 for SABMAR, while the temporal overlap between European rabbit and Egyptian mongoose was sensibly lower (Δ_1_ = 0.49 for SABOJI and Δ_1_ = 0.37 for SABMAR) (Fig 5).

**Figure 5.**
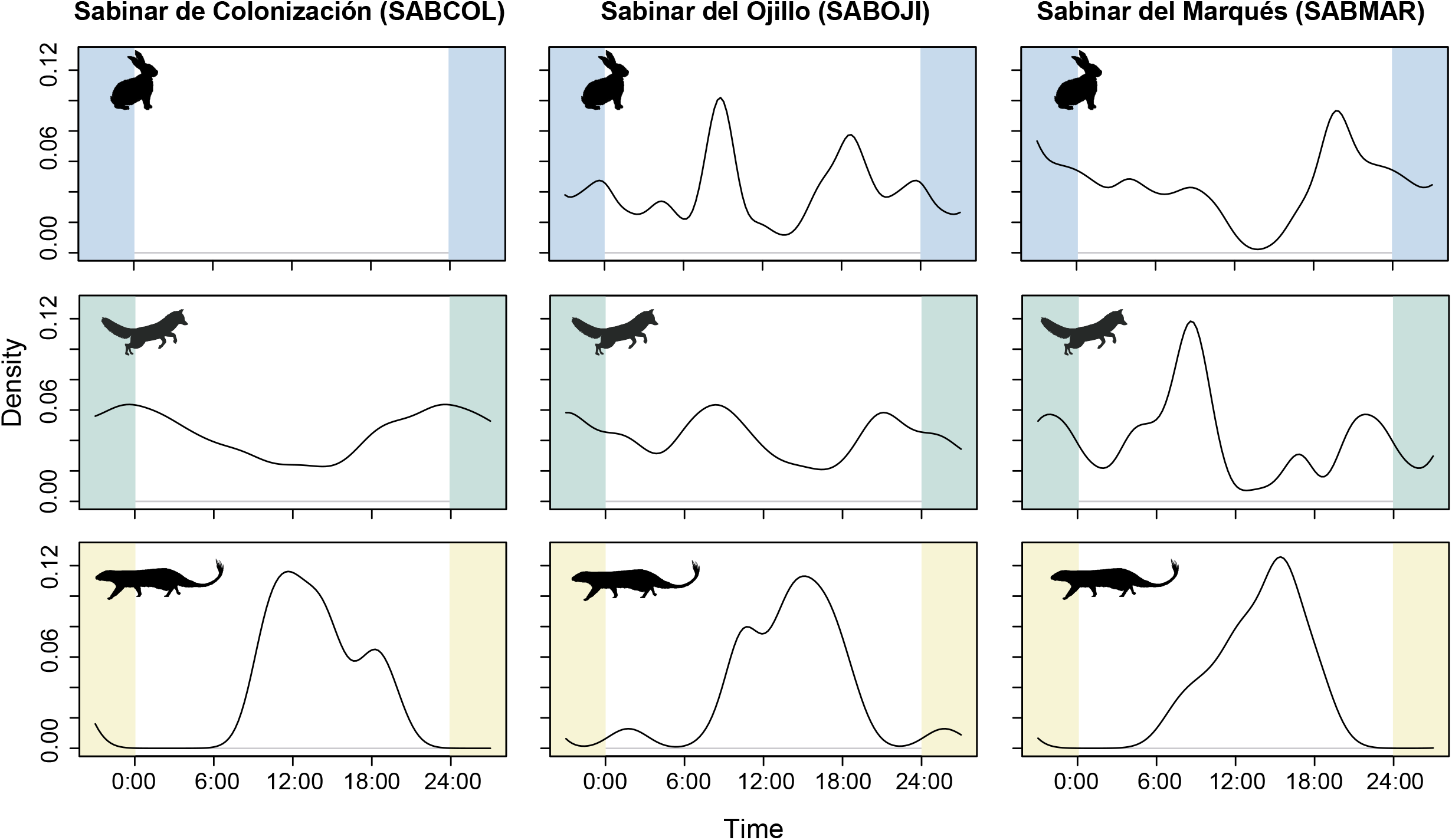
Diel activity patterns of two mesocarnivores (red fox and Egyptian mongoose; mid and bottom rows, respectively) and its main prey item (European rabbit, top row) along the gradient of ecological succession, from younger juniper stands (SABCOL) to mature, late successional stands (SABMAR).

### Temporal overlap and segregation between mesocarnivores

The average overlap and standard deviation of the activity patterns of the different mesocarnivores was Δ_1_ = 0.48 and SD = 0.29, which reflect a low, although variable, temporal overlap among these potential competitors. The average overlap values (Δ_1_ ± SD) of a species with respect to the set of potential competitors were: 0.59 ± 0.15 for the fox; 0.57 ± 0.34 for the badger; 0.50 ± 0.36 for the genet and 0.24 ± 0.18 for the Egyptian mongoose.

Strictly nocturnal species (European badger and common genet) showed a high overlap coefficient and no significant differences between their activity patterns (Δ_1_ = 0.79; W = 1.89, p-value = 0.386). In contrast, the Egyptian mongoose showed a strong and significant temporal segregation with the European badger (Δ_1_ = 0.16; W = 41.75, p-value < 0.01) and the common genet (Δ_1_ = 0.08; W = 65.79, p-value < 0.01) (Fig. 6). Simultaneously, we found that the activity pattern of the red fox had intermediate levels of temporal segregation with the common genet (Δ_1_ = 0.57; W = 24.96; p-value < 0.01) and the European badger (Δ_1_ = 0.71; W = 8.44; p-value = 0.014) and a low overlap with the Egyptian mongoose (Δ_1_ = 0.44; W = 73.48; p-value < 0.01) (Fig. 6). Temporal overlap between the red fox and the Egyptian mongoose varied per site, being Δ_1_= 0.41 for SABCOL, Δ_1_ = 0.45 for SABOJI and Δ_1_ = 0.36 for SABMAR (Fig. 5).

**Figure 6.**
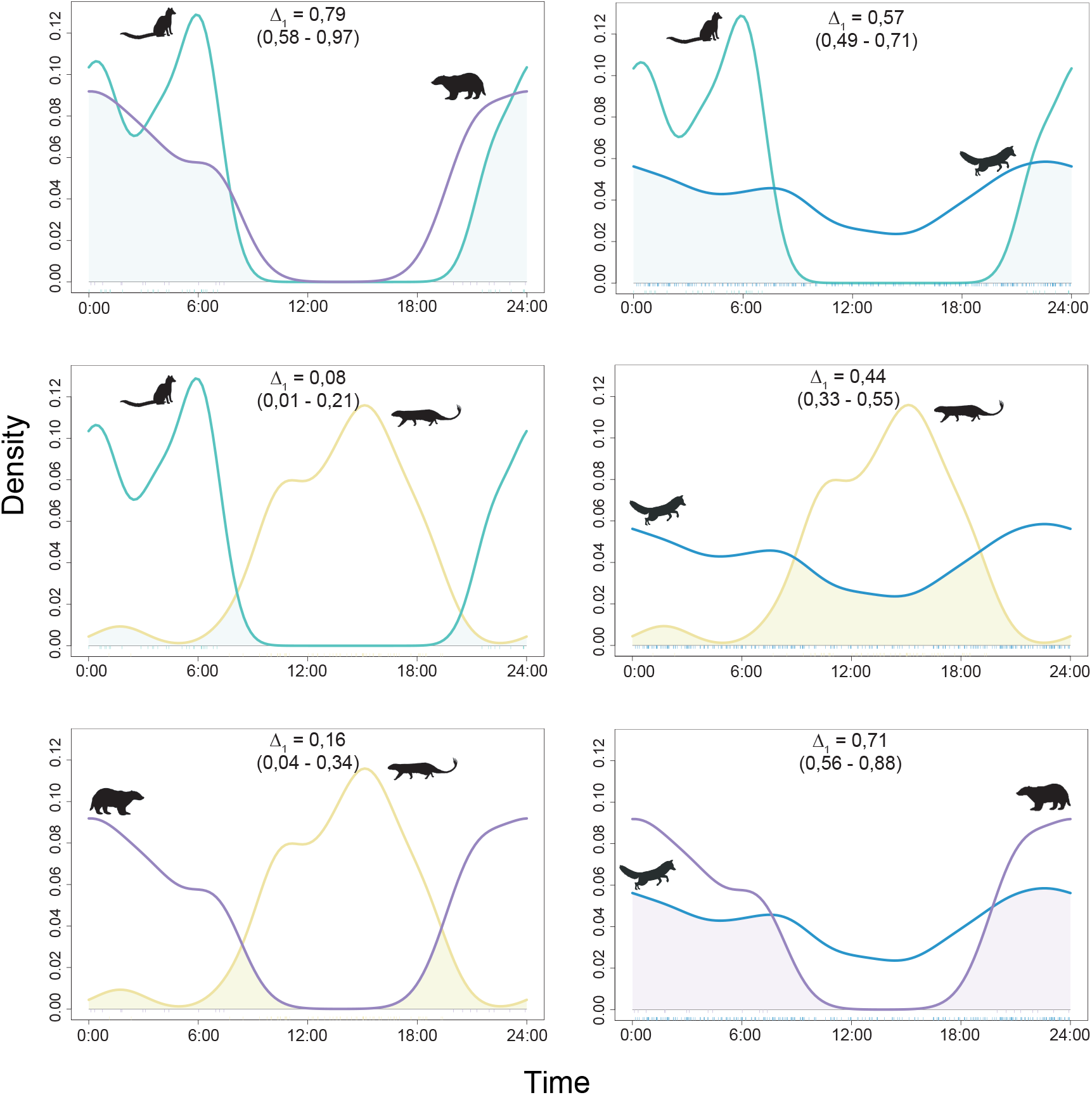
Kernel density plots and temporal overlap between mesocarnivores in the study area: common genet (green line), red fox (blue line), European badger (purple line) and Egyptian mongoose (yellow line). Overlap coefficients (Δ) and their respective 99% confidence intervals are shown in brackets.

### Activity centers and spatial overlap between predators

Predators showed differences in the location of their activity centers (peak RAI values) throughout the two years of sampling (Supplementary Fig. S2) and relatively low but variable spatial overlap values across the three study sites (Figure 7). On average, the red fox had the greatest spatial overlap with the rest of the mesocarnivores (ranging from PI (95% CI) = 0.27 (0.15-0.41) with European badgers to PI = 0.51 (0.36-0.66) with common genets); followed by the Egyptian mongoose (from PI = 0.17 (0.03-0-37) with European badgers to PI = 0.43 (0.35-0.56) with red foxes) and the common genets (from PI = 0.12 (0.03-0.27) with European badgers to PI = 0.51 (0.36-0.66) with red foxes) and lastly, the European badger (from PI = 0.12 (0.03-0.27) with common genets to PI = 0.27 (0.15-0.41) with red foxes) (Figure 7). In SABCOL, the highest values of spatial overlap correspond to the common genet and the red fox (PI = 0.54, 95% CI = 0.35-0.77) while the lowest values correspond to the European badger with the common genet and the Egyptian mongoose (PI = 0.05, 95%CI= 0-0.2; and PI = 0.02, 95% CI= 0-0.07, respectively). In SABOJI, the highest values correspond to the red fox and the Egyptian mongoose (PI = 0.70, 95% CI= 0.46-0.85) and the smallest value corresponds to the European badger and the Egyptian mongoose (PI = 0.03, 95% CI =0-0.13) while the rest of the pairwise comparisons showed intermediate values between 0.51 and 0.31. Finally, in SABMAR, the spatial overlap values are on average higher and more variable than in the other sites, ranging between the maximum value PI = 0.59 (95% CI =0.1-0.83; European badger and Egyptian mongoose) and the lowest PI = 0.19 (95% CI =0-0.52; common genet and Egyptian mongoose) (Supplementary Fig. S2).

**Figure 7.**
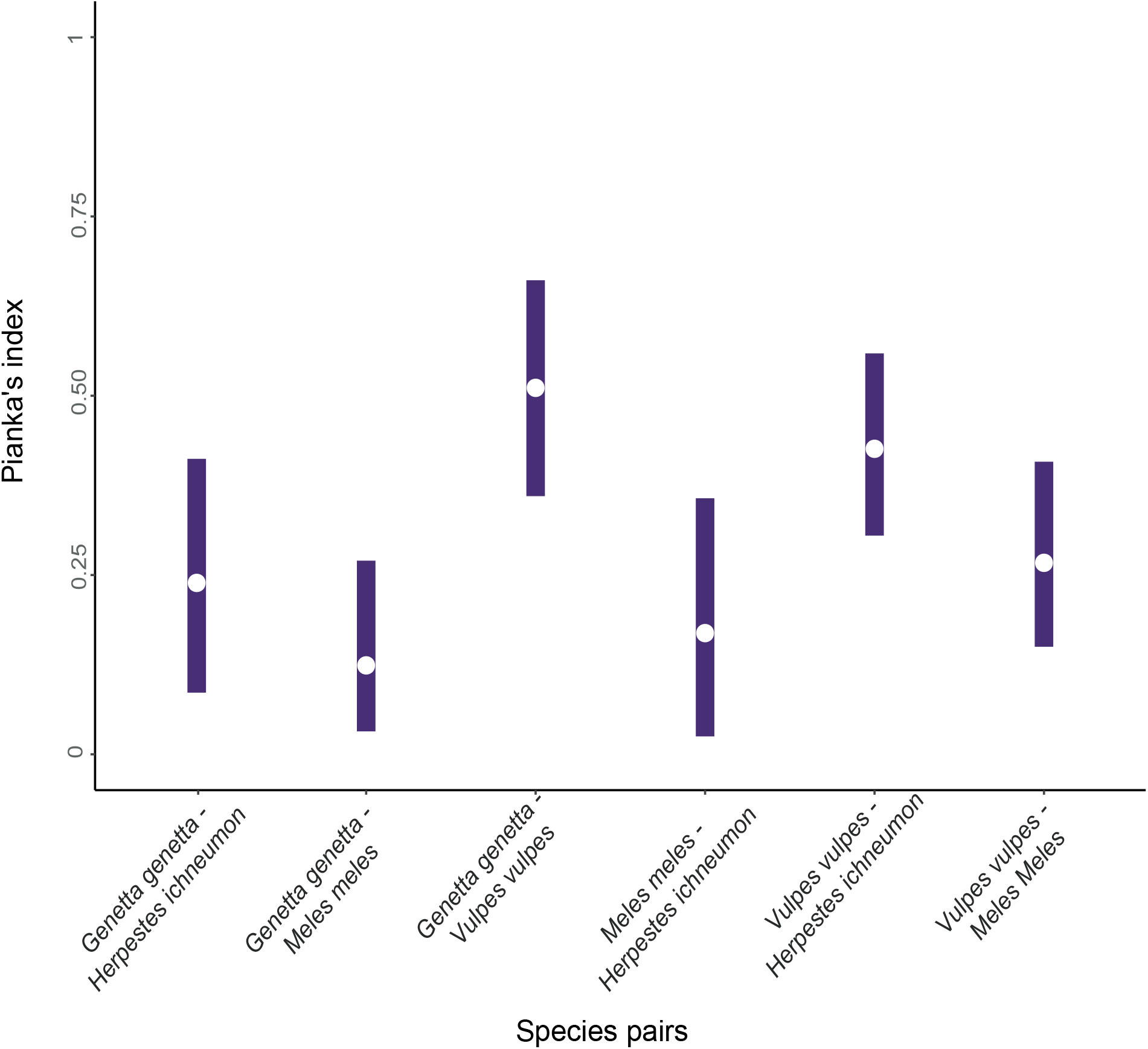
Spatial overlap for each pair of predators based on Pianka’s index (PI). White dots represent the estimated PI value and the bars indicate 95% confidence intervals.

## Discussion

In this study we explore the mechanisms underlying the structure of Mediterranean mesocarnivore communities living in sympatry in the absence of the top predator. We show that mesocarnivore coexistence is mediated by temporal, spatial, or trophic segregation when there is a greater overlap in any of these two niche axes. Unlike previous studies which have been traditionally focused on spatial responses over broad spatial scales (Fedriani et al. 1999, Soto and Palomares 2015, Monterroso et al. 2020, but see Barrull et al. 2014, Monterroso et al. 2014, Vilella et al. 2020), our study provides new insights on the small-scale coexistence of mesocarnivores released from top control, and reinforces the idea that the coexistence mechanisms underlying community structure are multidimensional. Overall, we found that temporal segregation facilitates co-occurrence of mesocarnivore species with similar trophic requirements (e.g., mongoose and genets), but that spatial segregation is discernible even at fine scale for species with high temporal and trophic overlap, with the larger mesocarnivores (fox and badgers) actively avoiding each other, putatively to not engage in IGK.

The highest values of temporal overlap occurred between the European badger and the common genet due to their predominantly nocturnal activity patterns. However, they showed lower levels of spatial overlap within their respective activity centers across study sites and marked differences in their respective trophic niches, mostly because of the preferential consumption of small mammals and birds by the common genet. Conversely, spatial overlap between genets and foxes was higher while they exhibit a moderate level of temporal and but also trophic segregation. According to our hypothesis, we found that the common genet has a narrower diet spectrum and showed higher levels of temporal overlap with its main prey (small mammals) whereas such synchrony does not occur with the activity patterns of the red-legged partridge. The common genet consumes mostly rodents and insectivores, with bird intake often considered as a secondary trophic resource (Palomares & Delibes, 1991a; Virgós et al., 1999). According to the Optimal Foraging Theory (MacArthur & Pianka, 1966), predators maximize their prey intake by showing an efficient foraging pattern. In the case of predators with narrower diets such as the genet, they would optimize their foraging behaviour by maximizing synchrony of their daily activity patterns to their main prey (Pyke et al., 1977), in this case, the small mammals.

In contrast, generalist predators may exhibit different activity patterns in response to differences in prey availability and their foraging strategies due to their eminently opportunistic behaviours. Interestingly enough, we found that the red fox showed a unimodal activity pattern with an activity peak around midnight in SABCOL, but shifted towards bimodality in sites where the rabbit was present, most likely to match the activity patterns of this prey and thus increase the probability of encounter. This was particularly obvious in SABOJI, where the temporal overlap between fox activity and that of the rabbits was the highest. Yet, this strategy was less beneficial in SABMAR, where an increase in the red fox activity around dawn may have triggered an anti-predatory behaviour in the European rabbit, which displayed only one activity peak at twilight, thus leading to lower temporal overlap.

However, temporal synchrony between badgers and mongooses with this prey item was on average relatively lower. Moreover, we did not find differences between the activity pattern exhibited by the Egyptian mongoose across study sites, and badger’s activity only occurred during night time in a consistent way across the different study sites. Dissimilarities in the synchrony of the activity patterns of the red fox and the Egyptian mongoose and European badgers with the most profitable prey in Doñana ((Ferreras et al., 2011; Martín et al., 1995; Palomares & Delibes, 1991b; Fedriani, 1996), the European rabbit, may be explained by their different foraging and consumption strategies of this food item. Egyptian mongooses and European badgers are species that preferentially consume younger rabbits by digging in burrows (Martín et al., 1995; Palomares & Delibes, 1991b). Conversely, the red fox resorts to a foraging strategy based mainly on the pursuit of prey items, although it also excavates in rabbit burrows (Fedriani et al., 1999). Thus, these distinct foraging strategies could explain why the Egyptian mongoose and European badgers display temporal asynchrony with the prey’s activity patterns while the red fox is expected to track and match the activity pattern in areas where this profitable prey item occurs.

Red foxes, European badgers and Egyptian mongooses are generalist predators known for their opportunistic feeding habits (Fedriani et al., 1999; Palomares, 1993; Rau, 1987). Yet, we found trophic segregation of the Egyptian mongoose with respect to the red fox and the European badger, with plant material (*e.g*., fleshy fruits) virtually absent from its diet. Meanwhile, the trophic overlap between European badger and red fox was the highest within the mesocarnivore community due to their omnivorous behaviour based on the consumption of fleshy fruits, invertebrates and lagomorphs. Interestingly, these three predators showed temporal and/or spatial segregation at microsite scale. Segregation along both axes is particularly remarkable for Egyptian mongooses and European badgers whose values of temporal and spatial overlap are considerably low. However, European badgers sustain higher levels of temporal overlap with red foxes, being the spatial avoidance at fine-scale the main mechanism to reduce the probability of interspecific encounters. Subtle differences in spatio-temporal dynamics while exploiting similar resources can reduce the probability of potentially adverse antagonistic encounters and promote coexistence between the most generalist predators within Doñana mesocarnivore guild.

Our results provide new insights into the traditional perspective regarding the mechanisms that allow coexistence between these four mesocarnivores. Previously, Soto & Palomares (2015) have shown that in relatively homogeneous habitats of Doñana, generalist mesocarnivores do not show clear segregation in habitat use on a relatively small spatial scale (69 2 x 2 km cells). However, we have shown how these generalist species, even coexisting in the same type of habitat and within sites of 1ha of surface, can show mechanisms of spatial avoidance at the microsite scale. These facts, together with the differences between activity patterns and trophic preferences, suggest that coexistence in this guild of mesocarnivores can be achieved through subtle differences in their respective ecological niches, with temporal segregation being the most pronounced mechanism among the different ecological niche dimensions studied here, but with small-scale spatial segregation diminishing the probability of antagonistic interspecific encounters.

Hence, our study addresses for the first time the interspecific relationships among the four mesocarnivores species present in Doñana National Park by integrating together their daily activity patterns, the use of the space, and their respective overlap, as well their trophic ecologies. Yet, these results should be taken with caution as the observational nature of this study does not allow us to establish a clear causal relationship in the observed patterns of the different species (e.g., temporal avoidance between competitors). Future research should address variations in the relative abundance of mesopredators across a larger number of sites, including several sites where rabbits are absent, and several sites where rabbits are present, with ideally the same mesocarnivore community across all sites. Unfortunately, our sample size (three sites and a low number of detections for certain species) was limited to address these questions and we recommend complementing camera trap data with other census methods (i.e., pellet counts or track surveys). Complementarily, trophic relationships between potential competitors could be evaluated by studying the diet composition through the collection, identification and analysis of faecal samples in the different study sites.

## Conclusions

The activity patterns of the different mesopredators are ultimately restricted by the circadian boundaries, but at the same time can be slightly modulated by interspecific interactions with both prey and potential competitors. Our results show that the coexistence of the different species that compose the mesocarnivore assemblage in the Doñana National Park can be facilitated by the temporal segregation among generalist species due to a divergence in the activity patterns (nocturnal, diurnal and cathemeral) but also by differences in trophic preferences among specialist species. Fine spatial scale monitoring of mesocarnivores activity patterns using camera traps for extended periods also revealed that local spatial segregation may result in low probabilities of interspecific encounter in these areas of close sympatry.

Our study further suggests that in order to clarify the coexistence mechanisms underlying community structure, it is necessary to consider a multidimensional perspective, addressing simultaneously multiple axes of the ecological niche. In markedly seasonal habitats like Doñana, with patchy vegetation including ample gradients of vegetation cover and successional status, even subtle differences in daily activity such as those documented here represent multiple pathways to facilitate sympatry of mesopredator species. These mechanisms might prove crucial for coexistence in settings where the apex predator is absent and medium-sized carnivores live at high densities and show ample overlap in resource use. Recent scenarios of rewilding with top predators may represent ideal conditions to experimentally examine the effects of mesopredator release in diverse carnivore guilds like those of the Mediterranean habitats studied here.

## Acknowledgements

We thank J.P. González-Varo, T. Burgos, E. Quintero, C. Gutierrez, R. Tarifa and J.L. Molina for technical support regarding species identification. We also thank P. Ferreras for sharing his trophic data with us. I. Mendoza and B. Arroyo gave helpful comments on early versions of the manuscript. Our work was funded by the project DISPMULTLAYER (CGL2017-82847-P), Spanish Ministry of Science and Innovation (PJ). JI was supported by the FPI grant PRE2018-085916 from the Spanish Ministry of Science and Innovation. ABL was supported by a Juan de la Cierva-Incorporación grant (IJCI-2017-31419), Spanish Ministry of Science, Innovation and Universities. We appreciate the logistic support from ICTS-RBD facilities during fieldwork.

## Author contributions

ABL conceived the original idea; IF collected the diet data, performed the analyses, designed the figures and wrote the first draft of the manuscript; JI and PJ designed the methodology; JI and IF compiled the camera-trap data. All authors contributed to the writing of the manuscript.

## Data accessibility

Data and R codes used in this study are deposited in Dryad (DOI: 10.5061/dryad.931zcrjkn) and GitHub (https://github.com/IagoFerreiroArias/TempPart).

## References

Agostinelli, C., & Lund, U. (2017). {R} package \texttt{circular}: Circular Statistics (version 0.4-93). CA: Department of Environmental Sciences, Informatics and Statistics, Ca’ Foscari University, Venice, Italy. UL: Department of Statistics, California Polytechnic State University, San Luis Obispo, California, USA. Retrieved from https://r-forge.r-project.org/projects/circular/

Arias-Del Razo, I., Hernández, L., Laundré, J. W., & Myers, O. (2011). Do predator and prey foraging activity patterns match? A study of coyotes (*Canis latrans*), and lagomorphs (*Lepus californicus* and *Sylvilagus audobonii*). Journal of Arid Environments, 75(2), 112–118.

Barrientos, R., & Virgós, E. (2006). Reduction of potential food interference in two sympatric carnivores by sequential use of shared resources. Acta Oecologica, 30(1), 107–116.

Barrull, J., Mate, I., Ruiz-Olmo, J., Casanovas, J. G., Gosàlbez, J., & Salicrú, M. (2014). Factors and mechanisms that explain coexistence in a Mediterranean carnivore assemblage: an integrated study based on camera trapping and diet. Mammalian Biology, 79(2), 123–131.

Batschelet, E. (1981). Circular statistics in biology. New York: Academic Press.

Bennie, J. J., Duffy, J. P., Inger, R., & Gaston, K. J. (2014). Biogeography of time partitioning in mammals. Proceedings of the National Academy of Sciences of the United States of America, 111(38), 13727–13732.

Chesson, P. (2000). Mechanisms of maintenance of species diversity. Annual Review of Ecology and Systematics, 31(1), 343–366.

Davis, M. L., Kelly, M. J., & Stauffer, D. F. (2011). Carnivore co existence and habitat use in the Mountain Pine Ridge Forest Reserve, Belize. Animal Conservation, 14(1), 56–65.

Di Bitetti, M. S., Di Blanco, Y. E., Pereira, J. A., Paviolo, A., & Pírez, I. J. (2009). Time Partitioning Favors the Coexistence of Sympatric Crab-Eating Foxes (*Cerdocyon thous*) and Pampas Foxes (*Lycalopex gymnocercus*). Journal of Mammalogy, 90(2), 479–490.

Díaz□Ruiz, F., Caro, J., Delibes Mateos, M., Arroyo, B., & Ferreras, P. (2016). Drivers of red fox (*Vulpes vulpes*) daily activity: prey availability, human disturbance or habitat structure? Journal of Zoology, 298(2), 128–138.

Ensing, E. P., Ciuti, S., De Wijs, F. A. L. M., Lentferink, D. H., Hoedt, A., Boyce, M. S., & Hut, R. A. (2014). GPS based daily activity patterns in european red deer and North American elk (*Cervus elaphus*): Indication for a weak circadian clock in ungulates. PLoS ONE, 9(9), e106997.

Fedriani, J. M. (1996). Dieta anual del zorro, *Vulpes vulpes*, en dos hábitats del Parque Nacional de Doñana. Doñana Acta Vertebrata, 23(2), 143–152.

Fedriani, J. M., Ferreras, P., & Delibes, M. (1998). Dietary response of the Eurasian badger, *Meles meles*, to a decline of its main prey in the Doñana National Park. Journal of Zoology, 245(2), 214–128.

Fedriani, J. M., Palomares, F., & Delibes, M. (1999). Niche relations among three sympatric Mediterranean carnivores. Oecologia, 121(1), 138–148.

Fenn, M. G. P., & Macdonald, D. W. (1995). Use of middens by red foxes: risk reverses rhythms of rats. Journal of Mammalogy, 76(1), 130–136.

Ferreras, P., Travaini, A., Zapata, S. C., & Delibes, M. (2011). Short-term responses of mammalian carnivores to a sudden collapse of rabbits in Mediterranean Spain. Basic and Applied Ecology, 12(2), 116–124.

Fisher, N. I. (1995). Statistical analysis of circular data. Cambridge University Press.

Foster, V. C., Sarmento, P., Sollmann, R., Tôrres, N., Jácomo, A. T. A., Negrões, N., … Silveira, L. (2013). Jaguar and puma activity patterns and predator□prey interactions in four Brazilian biomes. Biotropica, 45(3), 373–379.

Gaynor, K. M., Hojnowski, C. E., Carter, N. H., & Brashares, J. S. (2018). The influence of human disturbance on wildlife nocturnality. Science, 360(6394), 1232–1235.

Holt, R. D., Grover, J., & Tilman, D. (1994). Simple rules for interspecific dominance in systems with exploitative and apparent competition. The American Naturalist, 144(5), 741–771.

Jiménez, J., Nuñez-Arjona, J. C., Mougeot, F., Ferreras, P., González, L. M., García-Domínguez, F., … López-Bao, J. V. (2019). Restoring apex predators can reduce mesopredator abundances. Biological Conservation, 238, 108234.

Kahle, D. J., & Wickham, H. (2013). ggmap: spatial visualization with ggplot2. The R Journal, 5(1), 144–161.

Kronfeld-Schor, N., & Dayan, T. (2003). Partitioning of time as an ecological resource. Annual Review of Ecology, Evolution, and Systematics, 34(1), 153–181.

Lê, S., Josse, J., & Husson, F. (2008). {FactoMineR}: A Package for Multivariate Analysis. Journal of Statistical Software, 25(1), 1–18.

Linkie, M., & Ridout, M. S. (2011). Assessing tiger–prey interactions in Sumatran rainforests. Journal of Zoology, 284(3), 224–229.

López-Parra, M., Fernández, L., Ruiz, G., Gil-Sánchez, J. M., Simón, M. A., López, G., & Sarmento, P. (2012). Change in demographic patterns of the Doñana Iberian lynx *Lynx pardinus*: management implications and conservation perspectives. Oryx, 46(3), 403–413.

MacArthur, R. H., & Pianka, E. R. (1966). On optimal use of a patchy environment. The American Naturalist, 100(916), 603–609.

Martín, R., Rodríguez, A., & Delibes, M. (1995). Local feeding specialization by badgers (*Meles meles*) in a Mediterranean environment. Oecologia, 101(1), 45–50.

Monterroso, P., Alves, P. C., & Ferreras, P. (2013). Catch me if you can: Diel activity patterns of mammalian prey and predators. Ethology, 119(12), 1044–1056.

Monterroso, P., Alves, P. C., & Ferreras, P. (2014). Plasticity in circadian activity patterns of mesocarnivores in Southwestern Europe: Implications for species coexistence. Behavioral Ecology and Sociobiology, 68(9), 1403–1417.

Monterroso, P., Díaz□Ruíz, F., Lukacs, P. M., Alves, P. C., & Ferreras, P. (2020). Ecological traits and the spatial structure of competitive coexistence among carnivores. Ecology, 101(8), e03059.

Newsome, T. M., Greenville, A. C., Ćirović, D., Dickman, C. R., Johnson, C. N., Krofel, M., … Wirsing, A. J. (2017). Top predators constrain mesopredator distributions. Nature Communications, 8, 15469.

Palomares, F. (1993). Opportunistic feeding of the Egyptian mongoose, *Herpertes ichneumon* (L.) in Southwestern Spain. Revue d’écologie, 48, 295–304.

Palomares, F., & Caro, T. M. (1999). Interspecific killing among mammalian carnivores. The American Naturalist, 153(5), 492–508.

Palomares, F., & Delibes, M. (1991a). Alimentación del meloncillo *Herpestes ichneumon* y de la gineta *Genetta genetta* en la Reserva Biológica de Doñana, SO de la Península Ibérica. Doñana Acta Vertebrata, 18(1), 5–20.

Palomares, F., & Delibes, M. (1991b). Dieta del meloncillo (*Herpestes ichneumon*), en el Coto del Rey (Norte del Parque Nacional de Doñana, SO España). Doñana: Acta Vertebrata, 18(2), 197–194.

Palomares, F., & Delibes, M. (1991c). Ecología comparada de la gineta *Genetta genetta* (L.) y el meloncillo *Herpestes ichneumon* (L.) (Mammalia, Viverridae) en Doñana (SO de la Península Ibérica). Boletín de La Real Sociedad Española de Historia Natural (Sección Biología), 87(1–4), 257–266.

Palomares, F., & Delibes, M. (1994). Spatio-temporal ecology and behavior of European genets in southwestern Spain. Journal of Mammalogy, 75(3), 714–724.

Palomares, F., Ferreras, P., Fedriani, J. M., & Delibes, M. (1996). Spatial relationships between Iberian lynx and other carnivores in an area of south-western Spain. Journal of Applied Ecology, 33, 5–13.

Palomares, F., Ferreras, P., Travaini, A., & Delibes, M. (1998). Co-existence between Iberian lynx and Egyptian mongooses: estimating interaction strength by structural equation modelling and testing by an observational study. Journal of Animal Ecology, 67(6), 967–978.

Polis, G. A., Myers, C. A., & Holt, R. D. (1989). The ecology and evolution of intraguild predation: potential competitors that eat each other. Annual Review of Ecology and Systematics, 20(1), 297–330.

Pyke, G. H., Pulliam, H. R., & Charnov, E. L. (1977). Optimal foraging: a selective review of theory and tests. The Quarterly Review of Biology, 52(2), 137–154.

R Core Team. (2020). R: A Language and Environment for Statistical Computing. Vienna, Austria: R Foundation for Statistical Computing. Retrieved from https://www.r-project.org/

Rau, J. R. (1987). Ecología del zorro,” Vulpes vulpes (L.)”, en la reserva biológica de Doñana, SO de España. Universidad de Sevilla.

Revilla, E., & Palomares, F. (2002). Does local feeding specialization exist in Eurasian badgers? Canadian Journal of Zoology, 80(1), 83–93.

Ridout, M. S., & Linkie, M. (2009). Estimating overlap of daily activity patterns from camera trap data. Journal of Agricultural, Biological, and Environmental Statistics, 14(3), 322–337.

Rowcliffe, J. M., Kays, R., Kranstauber, B., Carbone, C., & Jansen, P. A. (2014). Quantifying levels of animal activity using camera trap data. Methods in Ecology and Evolution, 5(11), 1170–1179.

Rowcliffe, M. (2019). activity: Animal Activity Statistics. Retrieved from https://cran.r-project.org/package=activity

Santos, F., Carbone, C., Wearn, O. R., Rowcliffe, J. M., Espinosa, S., Lima, M. G. M., … Alvarez-Loayza, P. (2019). Prey availability and temporal partitioning modulate felid coexistence in Neotropical forests. PloS One, 14(3), e0213671.

Schoener, T. W. (1974). The compression hypothesis and temporal resource partitioning. Proceedings of the National Academy of Sciences of the United States of America, 71(10), 4169–4172.

Soto, C., & Palomares, F. (2015). Coexistence of sympatric carnivores in relatively homogeneous Mediterranean landscapes: functional importance of habitat segregation at the fine-scale level. Oecologia, 179(1), 223–235.

Soulé, M. E., Bolger, D. T., Alberts, A. C., Wrights, J., Sorice, M., & Hill, S. (1988). Reconstructed Dynamics of Rapid Extinctions of Chaparral□Requiring Birds in Urban Habitat Islands. Conservation Biology, 2(1), 75–92.

Valverde, J. A. (1957). Notes écologiques sur le lynx d’Espagne feus Lynx Pardina temminck. Terre et Vie (Revue d’Ecologie), 104, 51–67.

Valverde, J. A. (1967). Estructura de una comunidad mediterránea de vertebrados terrestres. Madrid: Consejo Superior de Investigaciones Científicas (España).

van Schaik, C. P., & Griffiths, M. (1996). Activity Periods of Indonesian Rain Forest Mammals. Biotropica, 28(1), 105–112.

Vazquez, C., Rowcliffe, J. M., Spoelstra, K., & Jansen, P. A. (2019). Comparing diel activity patterns of wildlife across latitudes and seasons: Time transformations using day length. Methods in Ecology and Evolution, 10(12), 2057–2066.

Vilella, M., Ferrandiz□Rovira, M., & Sayol, F. (2020). Coexistence of predators in time: Effects of season and prey availability on species activity within a Mediterranean carnivore guild. Ecology and Evolution, 00, 1–15.

Virgós, E., Llorente, M., & Cortésá, Y. (1999). Geographical variation in genet (*Genetta genetta* L.) diet: a literature review. Mammal Review, 29(2), 117–126.

Wang, Y., & Fisher, D. O. (2012). Dingoes affect activity of feral cats, but do not exclude them from the habitat of an endangered macropod. Wildlife Research, 39(7), 611–620.

Wickham, H. (2016). ggplot2: Elegant Graphics for Data Analysis. Springer-Verlag New York. Retrieved from https://ggplot2.tidyverse.org

Zhang, J., Ding, Q., & Huang, J. (2013). spaa: Species association analysis. R package version 0.2, 1.

Zielinski, W. J. (1986). Circadian rhythms of small carnivores and the effect of restricted feeding on daily activity. Physiology & Behavior, 38(5), 613–620.

