## Supplementary figures and images for "Fine-scale coexistence between Mediterranean mesocarnivores is mediated by spatial, temporal and trophic resource partitioning"

### Supplementary Fig. 1

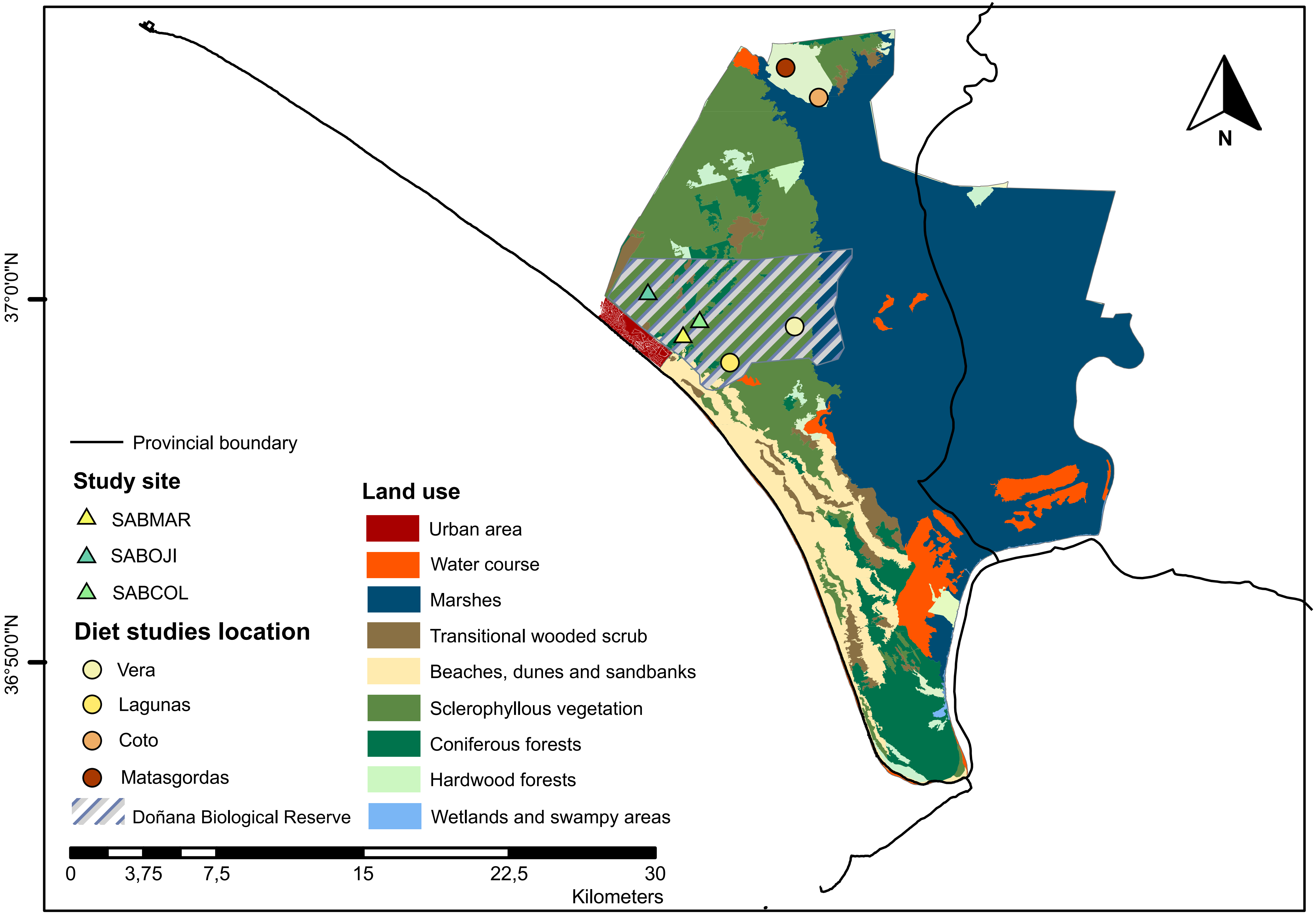


Distances from diet studies to study sites: Matasgordas, 14 km; Coto del Rey, 13,5km; Vera, 6.3km; and Lagunas, 3,7km.
