## Supplementary Table S1 for "Fine-scale coexistence between Mediterranean mesocarnivores is mediated by spatial, temporal and trophic resource partitioning"

**Table S1.** Summary of the studies used to carry out the correspondence analysis and explore the trophic relationships between mesocarnivores. Repetition of the studies is due to the different data coming from different areas, periods and/or species sampled within the same study.

| Study | Specie | Analysis | Area | Sample size | Habitat type | Feeding behaviour |
| --- | --- | --- | --- | --- | --- | --- |
| Fedriani (1996) | *Vulpes vulpes* | Scat analysis | Coto del Rey | 129 | Mediterranean scrubland | Generalist |
| Fedriani (1996) | *Vulpes vulpes* | Scat analysis | Coto del Rey | 164 | Dehesa | Generalist |
| Rau (1987) | *Vulpes vulpes* | Scat analysis | RBD | 781 | Pine forest, scrubland-marsh ecotone | Generalist |
| Ferreras et al. (2011) | *Vulpes vulpes* | Scat analysis | RBD  (1990) | 60 | Mediterranean  scrubland | Generalist |
| Ferreras et al. (2011) | *Vulpes vulpes* | Scat analysis | RBD  (1991-1992) | 112 | Mediterranean  scrubland | Generalist |
| Martín et al. (1995) | *Meles meles* | Scat analysis | RBD | 162 | Scrubland-marsh ecotone | Local specialization |
| Martín et al. (1995) | *Meles meles* | Scat analysis | RBD | 103 | Scrubland-marsh ecotone | Local specialization |
| Fedriani et al. (1998) | *Meles meles* | Scat analysis | RBD | 77 | Scrubland-marsh ecotone | Generalist |
| Fedriani et al. (1998) | *Meles meles* | Scat analysis | RBD | 68 | Scrubland-marsh ecotone | Generalist |
| Fedriani et al. (1998) | *Meles meles* | Scat analysis | Matasgordas | 108 | Scrubland-marsh ecotone | Generalist |
| Fedriani et al. (1998) | *Meles meles* | Scat analysis | Matasgordas | 101 | Scrubland-marsh ecotone | Generalist |
| Ferreras et al. (2011) | *Meles meles* | Scat analysis | RBD  (1990) | 69 | Mediterranean  scrubland | Generalist |
| Ferreras et al. (2011) | *Meles meles* | Scat analysis | RBD  (1991) | 72 | Mediterranean  scrubland | Generalist |
| Palomares (1993) | *Herpestes ichneumon* | Scat analysis | Vera | 190 | Marsh- scrubland ecotone | Generalist |
| Palomares (1993) | *Herpestes ichneumon* | Scat analysis | Lagunas | 60 | Dunes- scrubland ecotone | Generalist |
| Palomares (1993) | *Herpestes ichneumon* | Scat analysis | Coto del Rey | 178 | Mediterranean scrubland & pine forest | Local specialization |
| Ferreras et al. (2011) | *Herpestes ichneumon* | Scat analysis | RBD  (1990) | 41 | Mediterranean  scrubland | Generalist |
| Ferreras et al. (2011) | *Herpestes ichneumon* | Scat analysis | RBD  (1991) | 50 | Mediterranean  scrubland | Generalist |
| Ferreras et al. (2011) | *Herpestes ichneumon* | Scat analysis | RBD  (1992) | 13 | Mediterranean  scrubland | Generalist |
| Palomares and Delibes (1991a) | *Genetta genetta* | Scat analysis | RBD | 246 | Mediterranean scrubland | Specialist |
| Ferreras et al. (2011) | *Genetta genetta* | Scat analysis | RBD  (1990) | 72 | Mediterranean scrubland | Specialist |
| Ferreras et al. (2011) | *Genetta genetta* | Scat analysis | RBD  (1991) | 98 | Mediterranean scrubland | Specialist |
