## Supplementary Table S2 for "Fine-scale coexistence between Mediterranean mesocarnivores is mediated by spatial, temporal and trophic resource partitioning"

**Table S2.** Number of detections (N) of target species and trap success measured as number of detections of each species per 100 camera-days for each study site.

| Species | SABCOL | | SABOJI | | SABMAR | | Total |
| --- | --- | --- | --- | --- | --- | --- | --- |
|  | N | Trap success | N | Trap success | N | Trap success |  |
| Predators |  |  |  |  |  |  |  |
| *Vulpes. vulpes* | 165 | 5.98 | 72 | 2.56 | 64 | 2.07 | 301 |
| *Meles meles* | 6 | 0.22 | 5 | 0.18 | 11 | 0.36 | 22 |
| *Genetta genetta* | 26 | 0.94 | 9 | 0.32 | 7 | 0.23 | 42 |
| *Herpestes ichneumon* | 13 | 0.47 | 43 | 1.53 | 14 | 0.46 | 70 |
| Preys |  |  |  |  |  |  |  |
| *Oryctolagus cuniculus* | 6 | 0.22 | 340 | 12.0 | 304 | 10.0 | 650 |
| Small mammals | 15 | 0.54 | 30 | 1.07 | 53 | 1.74 | 98 |
| *Alectoris. rufa* | 42 | 1.52 | 3 | 0.11 | 2 | 0.065 | 47 |
