## Supplementary Table S3 for "Fine-scale coexistence between Mediterranean mesocarnivores is mediated by spatial, temporal and trophic resource partitioning"

**Table S3**. Number of detections of target species across three sites selected. Dawn: 1 hour before and after sunrise; Day: 1 hour after sunrise until 1 hour before sunset; Dusk: 1 hour before and after sunset; Night: 1 hour after sunset and 1 hour before sunrise. We set sunrise as 8:00:00 and sunset as 20:00:00. Rao Spacing Test significance: p-value <0,01 **; p value <0,05 *. Sampling effort per site reported as mean and standard deviation (*x̄* ± SD) of camera-days.

|  | SABCOL  (Sampling effort:80.3 ±28.3 days) | | | | |  | SABOJI  (Sampling effort: 80.4 ±27.3 days) | | | | |  | SABMAR  (Sampling effort: 86.8 ± 26 days) | | | | | Total | Rao Test (U) |
| --- | --- | --- | --- | --- | --- | --- | --- | --- | --- | --- | --- | --- | --- | --- | --- | --- | --- | --- | --- |
|  | Dawn | Day | Dusk | Night | Total |  | Dawn | Day | Dusk | Night | Total |  | Dawn | Day | Dusk | Night | Total |  |  |
| Predators |  |  |  |  |  |  |  |  |  |  |  |  |  |  |  |  |  |  |  |
| *V. vulpes* | 14 | 49 | 17 | 85 | 165 |  | 13 | 16 | 6 | 37 | 72 |  | 13 | 12 | 6 | 33 | 64 | 301 | 142.26* |
| *M. meles* | 0 | 0 | 3 | 3 | 6 |  | 1 | 0 | 0 | 4 | 5 |  | 0 | 0 | 1 | 10 | 11 | 22 | 178.95** |
| *G. genetta* | 1 | 0 | 0 | 25 | 26 |  | 1 | 0 | 0 | 8 | 9 |  | 0 | 0 | 0 | 7 | 7 | 42 | 165.81** |
| *H. ichneumon* | 0 | 13 | 0 | 0 | 13 |  | 1 | 38 | 2 | 2 | 43 |  | 1 | 13 | 0 | 0 | 14 | 70 | 199.78** |
| *Preys* |  |  |  |  |  |  |  |  |  |  |  |  |  |  |  |  |  |  |  |
| *O. cuniculus* | 3 | 2 | 0 | 1 | 6 |  | 71 | 96 | 45 | 128 | 340 |  | 27 | 82 | 49 | 146 | 304 | 650 | 156.97** |
| Small mammals | 2 | 0 | 0 | 13 | 15 |  | 0 | 1 | 3 | 19 | 23 |  | 4 | 0 | 9 | 47 | 60 | 98 | 198.27** |
| *A. rufa* | 1 | 41 | 0 | 0 | 42 |  | 0 | 3 | 0 | 0 | 3 |  | 0 | 2 | 0 | 0 | 2 | 47 | 226.53** |
